# Baculovirus-inducing fast-acting innate immunity kills *Plasmodium* liver stages

**DOI:** 10.1101/320036

**Authors:** Talha Bin Emran, Mitsuhiro Iyori, Yuki Ono, Fitri Amelia, Yenni Yusuf, Ashekul Islam, Asrar Alam, Ryohei Ogawa, Hiroyuki Matsuoka, Daisuke Yamamoto, Shigeto Yoshida

## Abstract

Baculovirus (BV), an enveloped insect virus with a circular double-stranded DNA genome, possesses unique characteristics that induce strong innate immune responses in mammalian cells. Here, we show that BV administration not only sterilely protects BALB/c mice for at least 7 days from subsequent *Plasmodium berghei* sporozoite infection but also eliminates existing liver-stage parasites completely, effects superior to those of primaquine, and does so in a TLR9-independent manner. Six hours post-BV administration, IFN-α and IFN-γ were robustly produced in serum, and RNA transcripts of interferon-stimulated genes were drastically upregulated in the liver. The *in vivo* passive transfer of post-BV administration serum effectively eliminated liver-stage parasites, and IFN-α neutralization abolished this effect, indicating that the BV liver-stage parasite killing mechanism is downstream of the type I IFN signaling pathway. Our results demonstrate that BV is a potent IFN-inducing prophylactic and therapeutic agent with great potential for further development as a new malaria vaccine and/or anti-hypnozoite drug.

## INTRODUCTION

Malaria remains a severe public health problem and causes significant economic losses worldwide. In 2016, there were approximately 216 million malaria cases and an estimated 445,000 malaria deaths, mainly in children under five (1). Malaria infection is initiated following injection of *Plasmodium* sporozoites into the skin during the taking of a blood meal by *Anopheles* mosquitoes. The sporozoites migrate to the liver and invade hepatocytes. Before clinical symptoms of malaria occur during the blood stage of infection, *Plasmodium falciparum* in the liver develop into exoerythrocytic schizonts for 5 to 6 days. *P. vivax* and *P. ovale* can develop dormant liver-stage forms, known as hypnozoites, which cause relapsing blood-stage infections months or years after the primary infection. Currently, the only licensed drug for the radical cure of *P. vivax* hypnozoites is primaquine (PQ), and artemisinin-based combination therapies are recommended by the World Health Organization (WHO) as the first-line treatment for blood-stage *P*. *falciparum* malaria. However, PQ has a high associated risk of life-threatening haemolytic anaemia in people with glucose-6-phosphate-dehydrogenase enzyme (G6PD) deficiency (2). For future malaria eradication strategies, safer radical curative compounds that efficiently kill hypnozoites are required.

A series of studies performed by Nussenzweig and colleagues in 1986–1987 revealed that exogenously administered interferon (IFN)-γ effectively inhibits development of liver-stage parasites *in vitro* and *in vivo* (3–6). Recently, Boonhok *et al*. reported that IFN-γ-mediated inhibition occurs at least partially in an autophagy-related protein-dependent manner in infected hepatocytes (7). Additionally, Liehl *et al*. reported that hepatocytes infected with liver-stage parasites induce a type I IFN secretion via the host cells sensing *Plasmodium* RNA, resulting in reduction of the liver-stage burden (8). These findings suggest that IFN-mediated immunotherapy against liver-stage parasites might be effective. However, new anti-hypnozoite drugs (e.g. recombinant IFNs or appropriate IFN inducers) have not been developed yet.

*Autographa californica* nucleopolyhedrosis virus (AcNPV), a type of baculovirus (BV), is an enveloped, double-stranded DNA virus that naturally infects insects. BVs possess unique characteristics that activate dendritic cell (DC)-mediated innate immunity through MyD88/Toll-like receptor 9 (TLR9)-dependent and -independent pathways (9). Takaku and colleagues reported that BV also directly activates murine natural killer (NK) cells through the TLR9 signalling pathway (10, 11), which leads to induction of NK cell-dependent anti-tumour immunity. Based on the unique adjuvant properties of BV that induce DC maturation and NK cell activation, which are prerequisites for generating robust and long-lasting adaptive immune responses, we have developed BV-based malaria vaccines effective for all three parasite stages, the pre-erythrocytic stage (12–14), asexual blood stage (15, 16), and sexual stage (17, 18).

Here, we investigated BV-mediated innate immunity against the pre-erythrocytic stage parasites. Our results clearly demonstrate that BV intramuscular administration not only elicits short-term sterile protection against *Plasmodium* sporozoite infection but also eliminates liver-stage parasites completely through the type I IFN signalling pathway. We propose that, due to its potent IFN-inducing function, BV has great potential for development into not only a new malaria vaccine platform capable of protecting vaccinators for a short period before and after malaria infection but also a new non-haemolytic single-dose alternative to PQ.

## RESULTS

### BV administration induce transgene expression and innate immune responses

This study investigated the effects induced by BV on innate immune responses relating to malaria infection. We used BES-GL3 harbouring two gene cassettes consisting of the *luciferase* gene under the control of the *CMV* promoter and the *DAF* gene under the control of the *p10* promoter, which were designed to express luciferase as a transducing marker and to display DAF as protection for BV from complement attack, respectively (19). BES-GL3 intramuscular administration into the left thigh muscle of mice initially increased the luciferase expression levels robustly, but these levels gradually decreased to 2% on day 28 (Fig. 1A), which is consistent with previous studies (19, 20). Among various cell types tested *in vitro*, hepatocytes were found to take up BV most effectively (21), suggesting a potential use for BV as a vector for liver-directed gene transfer. However, direct evidence of *in vivo* liver-directed gene transfer has not been reported because BV-mediated gene transfer into hepatocytes via intravenous injection is severely hampered by serum complement (22). We found for the first time that intravenous administration of BV effectively transduces hepatocytes *in vivo* (Fig. 1B).

**Figure 1.**
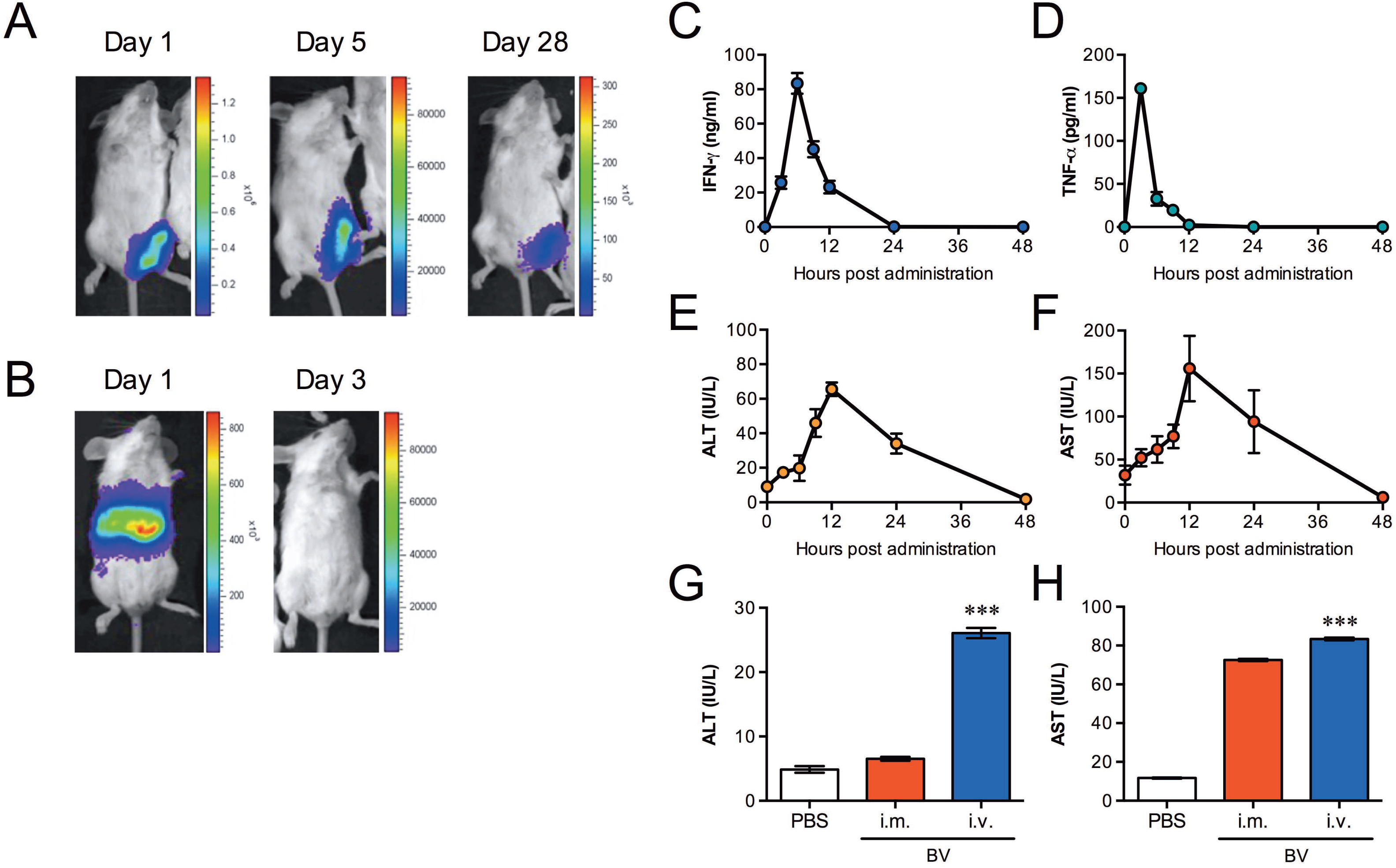
BV administration via intravenous vs intramuscular routes for transgene expression and innate immune responses. (A, B) Luciferase expression at different time points after intramuscular (10^8^ pfu) (A) or intravenous (10^7^ pfu) (B) administration of luciferase-expressing BES-GL3 (described as BV), detected by using the IVIS^®^ Lumina LT *in vivo* imaging system. The heat map image visible in the mice represents the total flux of photons (p/sec/cm2) in that area. Rainbow scales are expressed in radiance (p/s/cm^2^/sr). (C, D) Kinetics of proinflammatory cytokines, IFN-γ (C) and TNF-α (D), in the sera at different time points post-BV intravenous administration (10^7^ pfu) (n = 6). (E, F) Kinetics of liver damage markers. ALT (E) and AST (F) in the sera after BV intravenous administration (10^7^ pfu) (n = 6). (G, H) Comparison of ALT (G) and AST (H) in the sera at 6 h after intramuscular (10^8^ pfu) or intravenous (10^7^ pfu) administration of BV (n = 6). (C-H) Bars or points are the mean ± SD. (G, H) The difference from the PBS group was assessed by a Kruskal-Wallis test with Dunn’s correction. \*\*\**p* < 0.001. i.m., intramuscular; i.v., intravenous.

We next examined the kinetics of proinflammatory cytokines, ALT, and AST in sera following BES-GL3 intravenous administration. IFN-γ and TNF-α levels rapidly reached their peaks at 6 h and decreased to baseline by 24 h (Fig. 1, C and D). Similarly, the ALT and AST levels rapidly reached their peaks at 12 h and decreased to baseline by 48 h (Fig. 1, E and F). Compared with intravenous administration, intramuscular administration did not affect the ALT levels; although the AST level trended higher, this difference did not reach statistical significance (Fig. 1, G and H). ALT is a sensitive indicator of liver damage, so these results suggest that, for BV, intramuscular administration may be less destructive than intravenous administration.

### BV administration elicits sterile protection against sporozoite

Table 1 summarizes the protective efficacy results for BV administration against malaria sporozoite challenge. First, to examine the effects of BV intravenous administration, mice were intravenously administered 10^7^ pfu of BES-GL3. At 6 h post-BV injection, which coincides with peak IFN-γ production, the mice were intravenously challenged by 1,000 Pb-conGFP sporozoites, which are transgenic *P. berghei* constitutively expressing GFP. All BV-injected mice were protected, whereas all PBS- and AdHu5-injected mice treated similarly became infected. Next, we investigated the effects of BES-GL3 intramuscular administration (10^8^ pfu) followed by sporozoite challenge at various intervals post-BV injection. After intramuscular administration of BES-GL3, all mice were protected for at least 7 days. However, there was a complete loss of protection by 14 days post-BES-GL3 intramuscular administration, and no delay of parasitaemia was observed in these mice. Additionally, no protection was observed in mice treated intranasally with BES-GL3.

**Table 1.** Protective efficacy of BV injection 1 against *P. berghei* sporozoite challenge^a, b^

| Treatment<sup>a</sup> | Dose (Route<sup>c</sup>) | Time interval of challenge after administration | %Protection (protection/total)<sup>d</sup> |
| --- | --- | --- | --- |
| PBS | (i.v.) | 6 h | 0 (0/8)<sup>e</sup> |
| BV | 1 x 10<sup>7</sup> pfu (i.v.) | 6 h | 100 (8/8) |
| AdHu5-luc | 5 x 10<sup>7</sup> pfu (i.v.) | 6 h | 0 (0/3) |
| PBS | (i.m.) | 12 h – 14 days | 0 (0/20)<sup>e</sup> |
| BV | 1 x 10<sup>8</sup> pfu (i.m.) | 12 h | 100 (5/5) |
| BV | 1 x 10<sup>8</sup> pfu (i.m.) | 24 h | 100 (5/5) |
| BV | 1 x 10<sup>8</sup> pfu (i.m.) | 3 days | 100 (5/5) |
| BV | 1 x 10<sup>8</sup> pfu (i.m.) | 5 days | 100 (5/5) |
| BV | 1 x 10<sup>8</sup> pfu (i.m.) | 7 days | 100 (5/5) |
| BV | 1 x 10<sup>8</sup> pfu (i.m.) | 14 days | 0 (0/5) |
| BV | 1 x 10<sup>8</sup> pfu (i.n.) | 6 h | 0 (0/3) |
| BV | 1 x 10<sup>7</sup> pfu (i.v.) | 6 h/1,000 pRBC<sup>f</sup> | 0 (0/5) |
| CpG | 50 µg (i.m.) | 6 h | 90 (9/10)<sup>e</sup> |
| CpG | 50 µg (i.m.) | 24 h | 80 (4/5) |

<sup>a</sup>BALB/c mice were injected with BES-GL3 (described as BV) by the indicated route. After the indicated interval, mice were intravenously challenged with 1,000 Pb - conGFP sporozoites. Parasitaemia was monitored on days 5-8, 11, and 14 after sporozoite challenge. Once parasites appeared in the blood, all mice died.
<sup>b</sup>Scheme of the experimental design is shown in Fig. S2A.
<sup>c</sup>i.v., intravenous; i.n., intranasal; i.m., intramuscular
<sup>d</sup>Protection is defined as the complete absence of blood-stage parasitaemia on day 14 post-challenge.
<sup>e</sup>Cumulative data from two or four experiments <sup>f</sup>BALB/c mice were intravenously challenged with 1,000 Pb-conGFP-pRBC.

BES-GL3 intravenous administration failed to provide protection against challenge with 1,000 parasitized red blood cells (pRBCs) at 6 h post-BV injection, indicating that BV has no residual effect on blood-stage parasites. CpG intramuscular administration at 6 or 24 h prior to challenge conferred protection against sporozoite challenge in 90% or 80% of mice, respectively. This is consistent with previous work showing short-term (2-days) protection induced by CpG intramuscular administration (50 μg) against challenge with 100 *P. yoelii* sporozoites (23), although only partial protection (50%) was observed when the challenge occurred at 7 days post-CpG intramuscular injection. Thus, the protective efficacy induced by BES-GL3 intramuscular administration is more effective and longer-lasting (7 days) compared with CpG. All PBS-treated control mice developed blood-stage infection within 6 days following an intravenous injection of 1,000 Pb-conGFP sporozoites.

### BV administration completely eliminates of liver-stage parasites

Pathways stimulated by type I and II IFNs can lead to the killing of hepatocytes infected with liver-stage parasites (3–8). Because BV is a potent inducer of type I and II IFNs (24, 25), and we observed BV-mediated protection as described above, we next investigated whether BV-induced IFNs could kill liver-stage parasites *in vivo*. To examine the elimination effects on the trophozoite and exoerythrocytic (mature) schizont stages, we administered BES-GL3 intravenously or intramuscularly at two different intervals following sporozoite challenge, 24 and 42 h, respectively. Table 2 summarizes these results on the elimination efficacy of BES-GL3 administration against liver-stage parasites. Blood-stage parasites were completely prevented in all mice intravenously injected with BES-GL3 24 h post-infection; in contrast, the protective effectiveness of BES-GL3 intravenous administration was diminished when mice did not receive it until 42 h post-infection. The same results were obtained when mice were intramuscularly injected with BES-GL3.

**Table 2.** Elimination 1 of liver-stage parasites by BV administration^a, b^

| Treatment<sup>a</sup> | Dose (Route<sup>c</sup>) | Time interval of administration after challenge | %Elimination (uninfected/total) |
| --- | --- | --- | --- |
| PBS | (i.v.) | 24 h | 0 (0/12)<sup>e</sup> |
| BV | 1 x 10<sup>7</sup> pfu (i.v.) | 24 h | 100 (13/13)<sup>e</sup> |
| BV | 1 x 10<sup>7</sup> pfu (i.v.) | 42 h | 0 (0/3) |
| PBS | (i.m.) | 24 h | 0 (0/9)<sup>e</sup> |
| BV | 1 x 10<sup>8</sup> pfu (i.m.) | 24 h | 100 (7/7) |
| BV | 1 x 10<sup>6</sup> pfu (i.m.) | 24 h | 0 (0/5)<sup>f</sup> |
| BV | 1 x 10<sup>4</sup> pfu (i.m.) | 24 h | 0 (0/5)<sup>f</sup> |
| BV | 1 x 10<sup>8</sup> pfu (i.m.) | 42 h | 0 (0/3)<sup>f</sup> |
| PQ (High)<sup>d</sup> | 2 mg (i.p.) | 24 h | 100 (5/5) |
| PQ (Low)<sup>d</sup> | 0.1 mg (i.p.) | 24 h | 0 (0/5)<sup>f</sup> |

<sup>a</sup>BALB/c mice were intravenously injected with 1,000 Pb-conGFP sporozoites. After the indicated interval, mice were administrated either with BES-GL3 (described as BV) or PQ. Parasitaemia was monitored on days 5-8, 11, and 14 after sporozoite injection. Once parasites appeared in the blood, all mice died.
<sup>b</sup>Scheme of the experimental design is shown in Fig. S2B.
<sup>c</sup>i.v., intravenous; i.m., intramuscular; i.p., intraperitoneal
<sup>d</sup>The two different doses of PQ were administrated to eliminate liver-stage parasites. High dose (2 mg/100 µl) and low dose (0.1 mg/100 µl) were 533 and 27 times, respectively, higher than WHO recommended dose for human per weight.
<sup>e</sup>Cumulative data from three experiments.

To visualize the parasite elimination by BV, mice were infected with Pb-Luc, which is transgenic *P. berghei* constitutively expressing luciferase, and then examined via IVIS; this is a highly sensitive method for detecting liver- and blood-stage parasites. Parasites were observed in the liver at both 24 h and 42 h post-infection (Fig. 2, A and B, respectively; left panels). BES-GL3 intramuscular administration into the left thigh muscle at 24 h post-infection completely eliminated the liver-stage parasites at 72 h post-infection, whereas the PBS control treatment failed to prevent the development of blood-stage parasites (Fig. 2A; right panel). Although BES-GL3 intramuscular administration into the right thigh muscle at 42 h post-infection also failed to prevent the development of blood-stage parasites (Fig. 2B; right panel), it caused a significant delay of parasitaemia (Fig. 2C). This result indicates that even for exoerythrocytic schizonts (42 h post-infection), the elimination effect of BV intramuscular administration was invoked in the liver within 2–6 h because the exoerythrocytic merozoites of *P. berghei* are released from infected hepatocytes into the blood stream 44–48 h after the liver stage (26). Lower doses (10^4^ and 10^6^ pfu) of BES-GL3 administered at 24 h post-infection failed to prevent blood-stage parasites. However, a significant delay of parasitaemia was observed for the dose of 10^6^ pfu of BES-GL3 (Fig. 2D), indicating that the elimination effect is dependent on the amount of BV that is intramuscularly administered.

**Figure 2.**
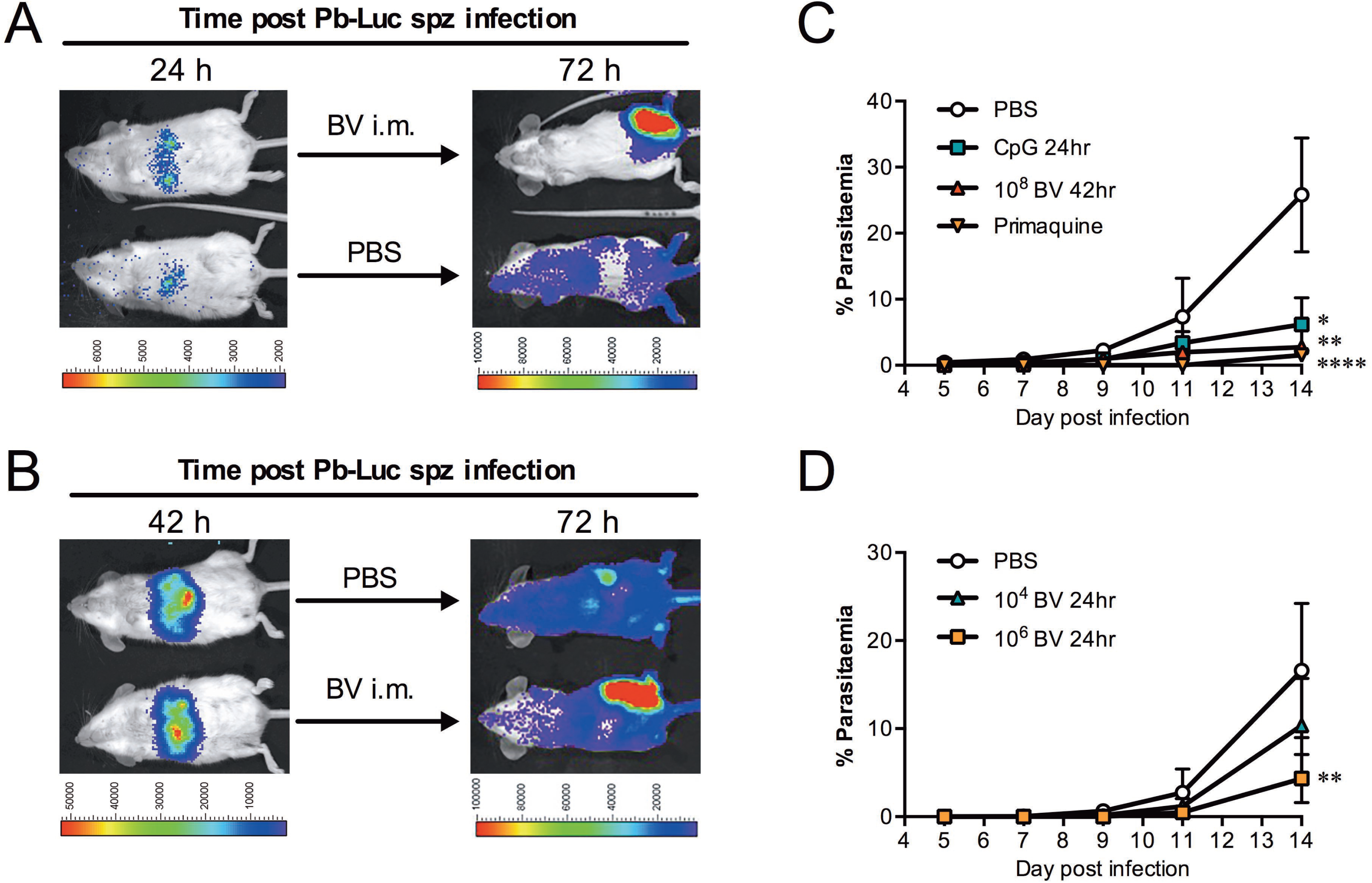
BV intramuscular administration completely eliminates liver-stage parasites. (A, B) Challenge infection with Pb-Luc sporozoites at 0 h, followed by BES-GL3 (described as BV) intramuscular administration (10^8^ pfu) at indicated time points. Luminescence in the liver shows parasite growth, whereas that in the thigh originates from the BV injection. The heat map images visible in the mice represent the total flux of photons (p/s/cm^2^) in that area. Rainbow scales are expressed in radiance (p/s/cm^2^/sr). (C, D) Delay of parasitaemia in infected mice. Parasitaemia of groups of infected mice shown in Table 2 (10^6^ or 10^4^ pfu of BV injected intramuscularly 24 h post-infection, 10^8^ pfu of BV injected intramuscularly 42 h post-infection, and PQ low dose administered 24 h post-infection) and Table 3 (CpG administered 24 h post-infection). Bars or points are the mean ± SD. The difference from the PBS group was assessed by a two-way ANOVA. \**p* < 0.05, \*\**p* < 0.01, \*\*\**p* < 0.001, \*\*\*\**p* < 0.0001. i.m., intramuscular; i.v., intravenous.

As PQ is the only licensed drug for the radical cure of *P. vivax* hypnozoites, we compared the elimination effects of BV with those of PQ. Two different doses of PQ, high dose (2 mg/mouse) and low dose (0.1 mg/mouse), were intraperitoneally administered; these doses are 533 and 27 times, respectively, higher than the WHO recommended dose per weight for people. A single administration of high dose PQ completely eliminated the liver-stage parasites (Table 2), whereas low dose PQ was suboptimal, producing only a reduction in parasite burden in the liver and significant delay of parasitaemia (Fig. 2C). The WHO-recommended treatment schedule for PQ is 15 mg/day for 14 days, but because high doses of PQ often cause side effects like nausea, vomiting, and stomach cramps, these side effects can limit patient compliance, potentially resulting in PQ resistance (27, 28). Thus, BV intramuscular administration may have important advantages of over PQ.

### BV-mediated liver-stage parasite killing is due to TLR9-independent pathways

CpG intramuscular administration completely eliminated early liver-stage parasites (6 h post-infection) (Table 3); however, although it caused a significant delay of parasitaemia, it had little effect on mature schizonts (24 h post-infection) (Fig. 2C). BV possesses unique characteristics that activate DC-mediated innate immunity through MyD88/TLR9-dependent and -independent pathways (9). Therefore, we next investigated whether TLR9 plays an important role in BV-mediated parasite killing in the liver. A single dose of intramuscularly administered BES-GL3 completely prevented blood-stage parasites in all TLR9^−/−^ mice previously infected with liver-stage parasites. In contrast, no elimination effect or parasitaemia delay was observed following CpG intramuscular administration in TLR9^−/−^ mice (Table 3). These results clearly demonstrate that BV-mediated parasite killing occurs via TLR9-independent pathways.

**Table 3.** Elimination of liver-1 stage parasites by BV injection in TLR9^−/−^ mice^a^

| Treatment<sup>a</sup> | Mouse strain | Dose | Time interval of administration after challenge | %Elimination (uninfected/total) |
| --- | --- | --- | --- | --- |
| PBS | TLR9<sup>-/-</sup> | - | 24 h | 0 (0/7) |
| BV | TLR9<sup>-/-</sup> | 1 x 10<sup>8</sup> pfu | 24 h | 100 (7/7) |
| BV | WT | 1 x 10<sup>8</sup> pfu | 24 h | 100 (5/5) |
| CpG | WT | 50 µg | 6 h | 100 (5/5) |
| CpG | WT | 50 µg | 24 h | 0 (0/4)<sup>b</sup> |
| CpG | TLR9<sup>-/-</sup> | 50 µg | 24 h | 0 (0/5) |

<sup>a</sup>TLR9<sup>-/-</sup> (BALB/c background) or WT mice were intravenously injected with 1,000 Pb-conGFP sporozoites. After 24 h, mice were intramuscularly administrated either with BES-GL3 (described as BV) or CpG ODN 1826 (described as CpG). Parasitaemia was monitored on days 5-8, 11, and 14 after sporozoite injection. Once parasites appeared in the blood, all mice died.
<sup>b</sup>Significant delay of parasitaemia was observed in infected mice, compared with the PBS group as showing in Fig. 2C.

BV intravenous administration was reported to produce type I IFNs through TLR-independent and IRF3-dependent pathways in mice (9). To further investigate IFN production following BV intramuscular administration, the IFN serum levels were measured in WT and TLR9^−/−^ mice 6 h after BES-GL3 intramuscular administration. As with intravenous administration, BES-GL3 intramuscular administration produced IFN-α in not only WT mice (6,311 ± 2,363 pg/ml) but also TLR9^−/−^ mice (1,590 ± 737 pg/ml), whilst mice intramuscularly injected with PBS or CpG did not produce detectable IFN-α (< 20.0 pg/ml) (Fig. 3A). IFN-γ, a type II IFN, was also produced in both WT mice (1,367 ± 1,303 pg/ml) and TLR9^−/−^ mice (488 ± 132 pg/ml) (Fig. 3B). CpG intramuscular administration induced much less IFN-γ compared with BV, but it induced a robust IL-12 response (Fig. 3C). Notably, CpG intravenous administration induced a high level of IFN-γ with considerable systemic side effects (29, 30).

**Figure 3.**
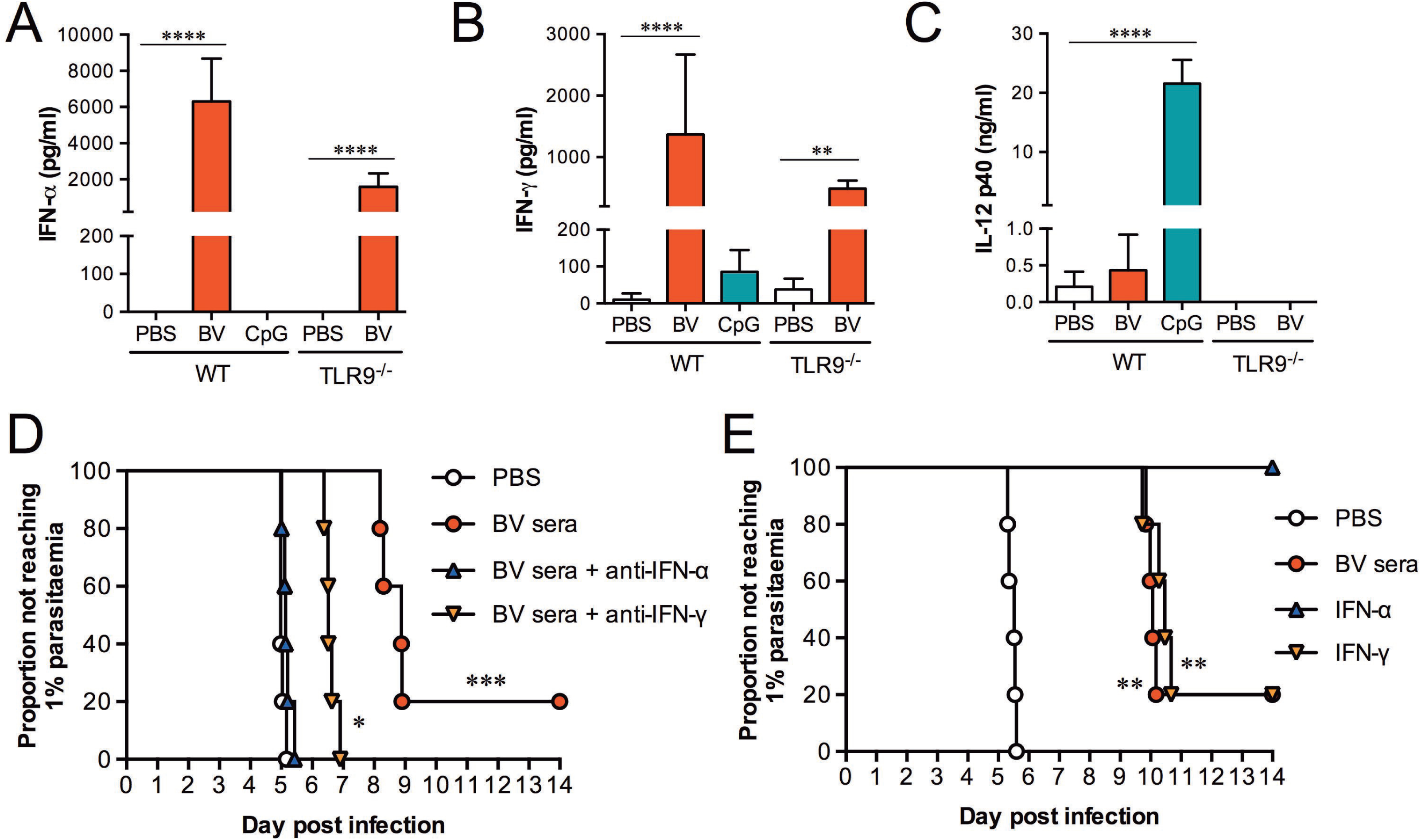
IFN-α induced by BV intramuscular administration contributes to elimination of liver-stage parasites. (A-C) Levels of IFN-α (A), IFN-γ (B), and IL-12 (C) in sera from WT or TLR9^−/−^ mice 6 h after intramuscular administration of BES-GL3 (described as BV) (10^8^ pfu), CpG, or PBS (n = 9−10). Bars are means ± SD. (D, E) Prediction of the time to reach 1% parasitaemia. (D) Results of serum transfer assay to determine the role of IFN-α and IFN-γ in the elimination of liver-stage parasites. Sera collected from mice 6 h after BV administration were neutralized by either anti-IFN-α or anti-IFN-γ antibody. Passive transfer of antibody-treated sera, non-treated sera, or PBS was conducted 24 h after sporozoite infection (n = 5). (E) Effect of exogenous IFN-α or IFN-γ on the elimination of liver-stage parasites. Recombinant mouse IFN-α or IFN-γ was intravenously administered 24 h after sporozoite infection (n = 5). (A-E) The difference from the PBS group was assessed by a Kruskal-Wallis test with Dunn’s correction. \**p* < 0.05, \*\**p* < 0.01, \*\*\**p* < 0.001, \*\*\*\**p* < 0.0001.

### Liver-stage parasites are killed by IFN-mediated immunity

To determine whether the serum cytokines act as effectors against liver-stage parasites, a serum transfer assay was performed. Pooled sera were collected from donor mice 6 h after they were intramuscularly injected with BES-GL3 or PBS. An aliquot of the pooled sera (100 μl/animal) was transferred to each recipient mouse 24 h after intravenous injection with 1,000 sporozoites. One of the five recipient mice effectively eliminated the liver-stage parasites, and the other four infected recipient mice showed a significant delay in the time to 1% parasitaemia (mean delay of 3.54 days; *p* = 0.0008, compared with the PBS sera group) (Fig. 3D).

We next examined whether neutralization of IFN-α or IFN-γ in the sera altered the effect of the sera on liver-stage parasites. Either anti-IFN-α or anti-IFN-γ antibody was incubated with 100 μl of the sera, which contained 8,619 pg/ml of IFN-α and 4,705 pg/ml of IFN-γ. Complete neutralization of IFN-α was confirmed by ELISA (see Fig. S1 in the supplemental material). The IFN-α- or IFN-γ-neutralized sera (100 μl) was intravenously administered to recipient mice that had been intravenously injected with 1,000 sporozoite 24 h before. The anti-IFN-α antibody treatment completely abrogated the serum-induced delay of parasitaemia, whereas the anti-IFN-γ antibody treatment only partially impaired the serum-induced elimination effect (Fig. 3D). To assess the effects of exogenous IFN-α and IFN-γ on elimination of liver-stage parasites, recombinant IFN-α (8,619 pg/mouse) or recombinant IFN-γ (4,705 pg/mouse) was intravenously administered to mice that had been intravenously injected with 1,000 sporozoites 24 h before. IFN-α administration completely eliminated the liver-stage parasites, whereas IFN-γ administration only partially eliminated them but caused a significant delay in the time to 1% parasitaemia (mean delay of 4.82 days; *p* = 0.0082, compared with the PBS group) (Fig. 3E). The IFN-α-mediated parasite elimination may be mediated via an effector mechanism distinct from that activated by IFN-γ. It is also possible that the effector mechanisms induced by IFN-α and IFN-γ may still be synergistically operative but that an alternate protective mechanism may be activated by BV. Similarly, Miller *et al*. showed that IFN-γ produced by NKT cells following type I IFN signalling from infected hepatocytes play an important role on elimination of liver-stage parasites (31). Table S1 summarizes the results on the elimination efficacy against liver-stage parasites of serum transfer and IFN administration.

### IFN-stimulated genes (ISGs) are upregulated in the liver post-BV intramuscular administration

Signal transduction of type I IFNs results in the induction of numerous ISGs (32). Some ISGs participate in direct antimicrobial activities, such as apoptosis induction and post-transcriptional event regulation for microbial killing, mainly acting as antiviral responses. Gene targeting studies have distinguished four effector pathways of the IFN-mediated antiviral response: the Mx GTPase pathway, 2’-5’ oligoadenylate-synthetase (OAS)-directed ribonuclease L pathway, protein kinase R (PKR) pathway, and ISG15 ubiquitin-like pathway (33). Additionally, several ISGs, such as IFN-induced proteins with tetratricopeptide repeats (IFITs), as well as the transcription factors IRF3 and IRF7 are responsible for sensing the liver-infection by *Plasmodium* sporozoites (8). To confirm the involvement of ISGs, the gene expression in the livers of mice intramuscularly injected with BES-GL3 were measured by quantitative RT-PCR (qRT-PCR). BES-GL3 significantly induced the gene expression of antiviral proteins *(Isg15, Mx1, Oas1a/b, Oasl1*, and *Pkr)* in wildtype (WT) mice (Fig. 4A). All these genes, except Oas1a/b, possibly due to the gene locus, were also upregulated by BV in TLR9^−/−^ mice. Gene expression of IFIT proteins, such as Ifit1, Ifit3, and Ifit44, were markedly and significantly enhanced in both WT and TLR9^−/−^ mice (Fig. 4B). Gene expression of the transcription factors Irf3 and Irf7 were also induced by BV in the same manner (Fig. 4C). These results indicate that systemic type I IFN secretion following BV intramuscular administration in the thigh muscle strongly induced ISGs in the liver.

**Figure 4.**
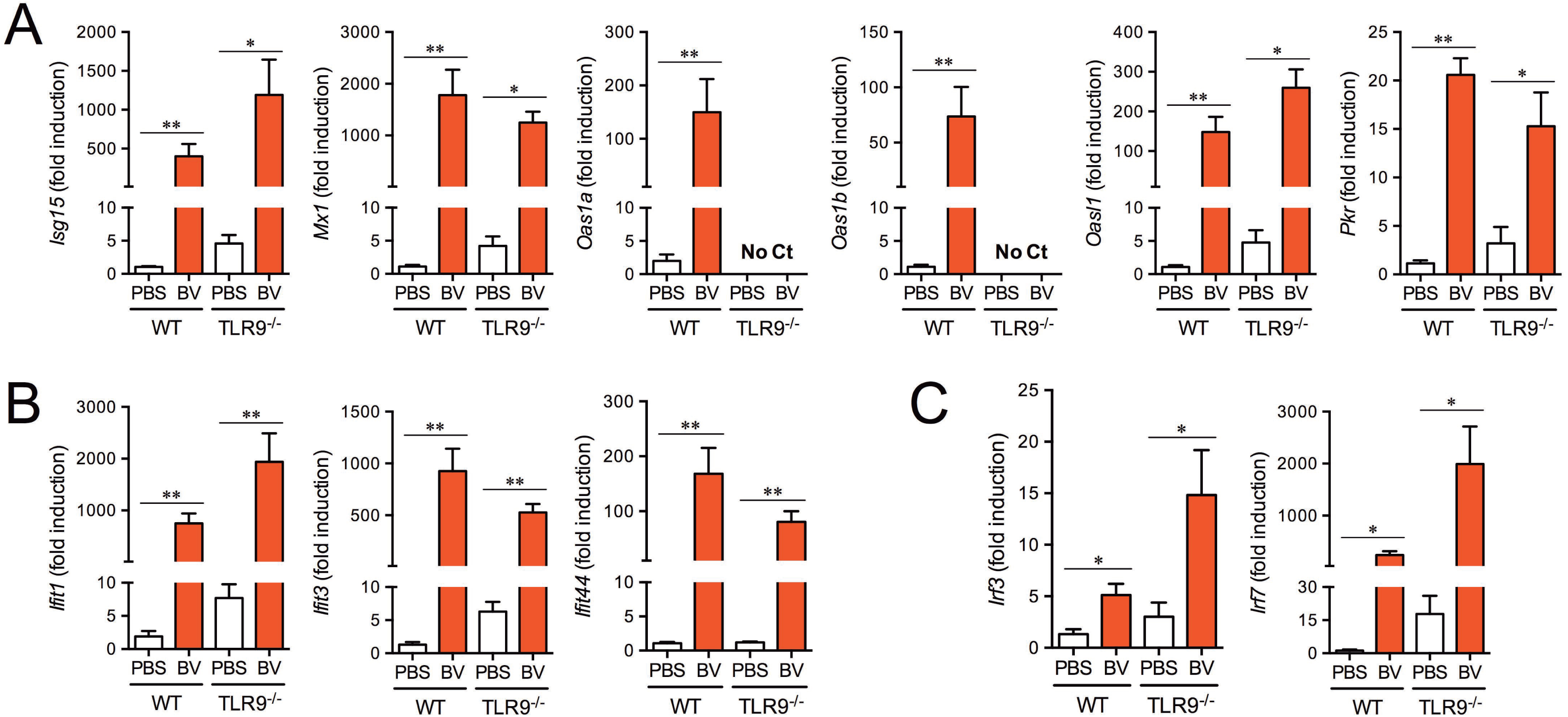
BV intramuscular administration induces ISGs in the liver. Gene expression of antiviral proteins (A), IFITs (B), and IRFs (C) in the livers of WT and TLR9^−/−^ mice at 6h after intramuscular administration of BES-GL3 (described as BV) (10^8^ pfu) was measured by real-time RT-PCR (n = 5−7). Bars are means ± SEM. The difference from the PBS group was assessed by a Mann-Whitney’s U test. \**p* < 0.05, \*\**p* < 0.01.

### AdHu5-prime/BDES-boost heterologous immunization regimen confers sterile protection and complete elimination

To evaluate our newly developed malaria vaccine in an AdHu5-prime/BDES-boost heterologous immunization regimen (14), mice were challenged twice (before and after) BDES-sPfCSP2-boost following AdHu5-prime. All mice survived without any symptoms following both the first and second challenges (Table S2, Group 2). In contrast, AdHu5-prime immunization alone did not confer protection (Group 1). All control mice intramuscularly injected with PBS became infected (Groups 3 and 4). Thus, BDES-PfCSP boosting was able to exert not only a therapeutic effect on liver-stage parasites but also a prophylactic effect on sporozoites. The animal experimental designs are illustrated in Fig. S2, E and F.

## DISCUSSION

Here, we show that BV intramuscular administration not only elicits short-term sterile protection against sporozoite infection but also eliminates liver-stage parasites completely. For liver-stage parasites proliferating vigorously at 24 h post-infection, the BV-induced fast-acting innate immune responses completely kill them within the following 20 h and prevent blood-stage parasite development in the absence of any clinical symptoms, which is more effective than PQ in a mouse model with early liver-stage *P. berghei*. The *P. berghei* liver-stage model is thought to correlate with anti-hypnozoite activity in primates (34). PQ is currently the only available drug that kills the dormant hypnozoites of *P. vivax*, but its severe side effects in G6PD-deficient people prevent the widespread use of this drug (35). The presence of hypnozoites and their drug-insensitivity form a major hurdle for malaria elimination programmes. Although BV has not been tested in a clinical setting yet, our previous study showed that the BV-based vaccine vector is safe and well-tolerated with acceptable reactogenicity and systemic toxicity in a primate model (13). Thus, BV offers a promising new non-haemolytic single-dose alternative to PQ for first-in-human clinical trials. Further experiments to determine optimum BV administration routes and dosages are needed.

BV possesses attractive attributes as a new vaccine vector (e.g., its low cytotoxicity, inability to replicate in mammalian cells and absence of pre-existing antibodies against it). This study demonstrates the further unique advantage of BV as a therapeutic vaccine vector with short-term protection via an intrinsic potent immunostimulatory property. In a Phase II–III malaria vaccine trial, all volunteers are presumptively treated with three daily doses of anti-malaria drug one week before final vaccination and rechecked for asexual *P. falciparum* parasitaemia one week after the final vaccination. Any subject who tests parasite-positive is treated with a second line drug or excluded from the trial (36). Thus, clinical trials aim to test vaccine efficacy after all vaccine schedules are completed to assess the maximum effect. For clinical application, however, vaccinators are still in danger of infection until the full vaccination schedule is completed, even though improved effective vaccine would be developed. This study shows that our newly developed heterologous AdHu5-PfCSP-prime and BDES-PfCSP-boost vaccine eliminated liver-stage parasites that had infected the mice 24 h before administration of the BDES-boost and also elicited sterile protection against sporozoite challenge 21 days post-boost. We propose that BV-based vaccines can not only minimize the risk of infection for vaccinators during the vaccination schedule but also generate robust and long-lasting adaptive immune responses via stimulation of the innate immune system. Alternatively, BV itself may also be used as an additive to eliminate liver-stage parasites and impart this short-term protection to RTS,S or other licensed vaccines.

This study showed that IFN-α and IFN-γ were rapidly and robustly produced in serum 6 h post-BV administration. Interestingly, the prophylactic effect against sporozoite infection lasted for at least 7 days, even after IFN levels returned to baseline. In the case of ‘natural’ *Plasmodium* liver-stage infection, the infected hepatocytes induce IFN-α, resulting in a reduction of the liver-stage burden (8). However, the parasite-induced IFN-α responses fail to eliminate every parasite. This implies that the endogenous innate immune responses may be not strong enough for complete elimination and/or that the innate immune response peak occurs at the end of liver-stage development, just prior to or concurrent with exoerythrocytic merozoite release (37). Compared with the type I IFN induced by host sensing of parasites, the quantity of BV-induced type I IFN and its speedy induction may make it more effective, resulting in its potent therapeutic and prophylactic effects. A better understanding of the molecular mechanisms by which BV administration confers both protection and elimination of pre-erythrocytic parasites will provide new strategies for malaria drug and vaccine development.

IFN-α has been extensively explored for its efficacy in various disease conditions and is currently used as a standard treatment in several illnesses. However, its use is accompanied by a wide variety of possible side effects (38), such as autoimmune thyroiditis. This study found that BV intramuscular administration, which induced 8,619 pg/ml of IFN-α in mouse sera with normal levels of ALT, completely killed liver-stage parasites. The manufacturing cost of BV would be much lower than that of recombinant IFN-α. Thus, BV intramuscular administration also has great potential for use as an alternative IFN-α-based immunotherapy; its high biological activity, cost-effectiveness, non-invasive nature, and minimal adverse effects make it superior to the current IFN-α therapy using recombinant IFN-α via intravenous administration.

In conclusion, BV effectively induces fast-acting innate immune responses that provide powerful first lines of both defensive and offensive attacks against pre-erythrocytic parasites. Our results illustrate the great potential of baculovirus as a new potent prophylactic and therapeutic immunostimulatory agent against preerythrocytic-stage parasites. We propose that these baculovirus-based vaccine and drug strategies could be applicable not only for malaria but also for other infectious diseases.

## MATERIALS AND METHODS

### Animals, cell lines, parasites, and mosquitoes

Female inbred BALB/c (*H-2^d^*) mice were obtained from Japan SLC (Hamamatsu, Shizuoka, Japan) and used in all experiments at 7–8 weeks of age. TLR9^−/−^ deficient (TLR9^−/−^) mice on a BALB/c background were kindly provided by S. Akira (University of Osaka, Suita, Japan). *Spodoptera frugiperda* and HepG2 cells were maintained as described previously (13). Three transgenic *P. berghei* ANKA parasites were used in this study: GFP-P *berghei* (Pb-conGFP) (39), luciferase-*P berghei* (Pb-Luc) (40), and PfCSP-*P berghei* (PfCSP-Tc/Pb) (41). These transgenic parasites were maintained by cyclical passaging through BALB/c mice and *Anopheles stephensi* (SDA 500 strain) at the Kanazawa University and Jichi Medical University according to a standard protocol (12, 42).

### Ethics statement

All animal care and handling procedures were approved by the Animal Care and Ethical Review Committee of Kanazawa University (no. 22118–1) and Jichi Medical University (no. 09193) Japan. For animal experiments, all efforts were made to minimize suffering in the animals. Mice were anesthetized with ketamine (100 mg/kg; intramuscular; Daiichi Sankyo, Tokyo, Japan) and xylazine (10 mg/kg; intramuscular; Bayer, Tokyo, Japan) when necessary.

### Recombinant viruses

The recombinant baculoviruses BES-GL3 and BDES-sPfCSP2-WPRE-Spider have been described previously (19). The purified baculovirus particles were free of endotoxin (<0.01 endotoxin units/10^9^ pfu), as determined by the Endospecy^®^ endotoxin measurement kit (Seikagaku Co., Tokyo, Japan). The recombinant adenoviruses AdHu5-Luc and AdHu5-sPfCSP2 have been described previously (14). In this paper, BDES-sPfCSP2-WPRE-Spider and AdHu5-sPfCSP2 are described as BDES-PfCSP and AdHu5-PfCSP, respectively.

### Collection of sporozoites

*An. stephensi* mosquitoes were infected by feeding on infected mice using standard methods of mosquito infection. On days 21–24 post-infection, the salivary glands of the mosquitoes were collected by hand-dissection. Salivary glands were collected in DMEM (Thermo Fisher Scientific K.K., Tokyo, Japan) and homogenized in a plastic homogenizer. The free sporozoites were counted in a disposable haemocytometer counting chamber using phase-contrast microscopy.

### Analysis of protective effects against sporozoite parasites

BALB/c mice were intravenously, intramuscularly, or intranasally administered 10^4^–10^8^ pfu of BES-GL3. Alternatively, instead of BES-GL3, BALB/c mice were intramuscularly injected with 50 μg of CpG ODN 1826 (TCCATgACgTTCCTgACgTT, Fasmac Inc., Tokyo, Japan). The mice were intravenously challenged with 1,000 Pb-conGFP sporozoites or 1,000 pRBCs at various time intervals (6 h–14 days). The mice were checked for *P*. *berghei* blood-stage infection by microscopic examination of Giemsa-stained thin smears of their tail blood, prepared on days 5, 6, 7, 8, 11, and 14 post-challenge. The time required to reach 1% parasitaemia was determined as described previously (43). A minimum of 20 fields (magnification, × 1,000) were examined before a mouse was deemed to be negative for infection. Protection was defined as the complete absence of blood-stage parasitaemia on day 14 post-challenge.

### Analysis of elimination effects on liver-stage parasites

BALB/c mice were intravenously injected with 1,000 Pb-conGFP sporozoites and subsequently intravenously (10^7^ pfu) or intramuscularly (10^8^ pfu) injected with BES-GL3 at various time intervals (6, 24, or 42 h post-infection). Alternatively, instead of BV, a single high (2 mg) or low (0.1 mg) dose of PQ (primaquine diphosphate 98%, Sigma-Aldrich, St. Louis, MO, USA), with corresponding concentrations of roughly 40 mg/kg body weight and 20 mg/kg body weight respectively, was intraperitoneally administered 24 h after injection of 1,000 Pb-conGFP sporozoites. The mice were checked for *P. berghei* blood-stage infection and evaluated for 1% parasitaemia as described above.

### *In vivo* bioluminescent imaging

BALB/c mice were intravenously or intramuscularly injected with BES-GL3 on day 0, and D-luciferin (15 mg/ml; OZ Biosciences, Marseille, France) was then intraperitoneally administered (150 μl/mouse) to these mice at various time points. The animals were anesthetized with a ketamine (100 mg/kg)/xylazine (10 mg/kg) mixture 10 min later, and the luciferase expression was detected with an IVIS^®^ Lumina LT *in vivo* imaging system (PerkinElmer, Waltham, MA, USA). Alternatively, BALB/c mice were intravenously injected with 1,000 Pb-Luc sporozoites followed by BES-GL3 (10^8^ pfu) intramuscular administration into the left thigh muscle 24 or 42 h later. At 72 h after the sporozoite injection, the luciferase expression was detected as described above. At days 5–14 post-infection, the same mice were analysed for blood-stage infections by determination of the course of parasitaemia in Giemsa-stained thin blood films of tail blood.

### Cytokine, AST, and ALT assays

BALB/c mice were intravenously or intramuscularly injected with BV, and sera were subsequently harvested from whole blood obtained by cardiopuncture at various times and stored at −20 °C until analysis. The concentrations of cytokines in the sera were determined by sandwich ELISA using a Mouse IFN-γ ELISA MAX™ standard kit (Biolegend Inc., San Diego, CA, USA), mouse IL-12/IL-23 (p40) ELISA MAX™ standard kit (Biolegend Inc.), or mouse TNF-α ELISA MAX™ deluxe kit (Biolegend Inc.) according to the manufacturer’s instructions. The concentration of IFN-α was determined by sandwich ELISA as described previously. In brief, rat monoclonal antibody against mouse IFN-a (PBL Biomedical Laboratories clone RMMA-1, Piscataway, NJ, USA) was used as the capture antibody (2 μg/ml for coating), rabbit polyclonal antibody against mouse IFN-α (PBL Biomedical Laboratories) was used at 80 neutralizing units per ml for detection, and HRP-conjugated goat anti-rabbit IgG (Bio-Rad, Hercules, CA, USA) was used as the secondary reagent. Recombinant mouse IFN-α (PBL Biomedical Laboratories) was used as the standard. The lower detection limits for the IFN-γ and the IFN-α immunoassays were each <20 pg/ml, whereas those for the IL-12 and the TNF-α immunoassays were each <10 pg/ml. The concentrations of ALT and AST in the sera were determined by using a GPT/GOT assay kit (Transaminase CII-test; Wako Pure Chemical Industries, Ltd., Tokyo, Japan) according to manufacturer’s instructions.

### Serum transfer and IFN administration analysis

Pooled sera were obtained from blood harvested by cardiopuncture from 5 BALB/c mice that had been intramuscularly injected with BES-GL3 6 h previously (at −6 h), and the concentrations of IFN-α and IFN-γ were measured immediately. On the same day, the IFN-α and IFN-γ in 100-μl aliquots of the pooled sera were neutralized by incubation with sufficient amounts of anti-IFN-α (anti-mouse interferon alpha, rabbit serum; PBL Biomedical Laboratories) and anti-IFN-γ (Ultra-LEAF™ purified anti-mouse IFN-γ antibody; BioLegend Inc.) antibodies, respectively, on ice for 6 h according to the manufacturer’s instructions. At 24 h after being intravenously injected with 1,000 Pb-conGFP sporozoites, BALB/c mice were then intravenously injected with 100 μl of the sera that had been treated with either anti-IFN-α or anti-IFN-γ. For the IFN administration experiment, BALB/c mice that had been intravenously injected with 1,000 Pb-conGFP sporozoites 24 h before were then intravenously administered either 8,619 pg of IFN-α or 4,705 pg of IFN-γ. For each experiment, the mice were checked for *P. berghei* blood-stage infection and evaluated for 1% parasitaemia as described above.

### RNA isolation from livers and qRT-PCR quantification

BALB/c (WT or TLR9^−/−^) mice were intramuscularly injected with 10^8^ pfu of BES-GL3. Alternatively, 50 μg of CpG ODN1826 were administered intramuscularly. Six hours later, whole livers were obtained by dissection of the treated mice. Each whole liver was placed in a 5-ml plastic tube with a cap containing 4 ml of buffer RLT (Qiagen, Valencia, CA, USA) containing 1% 2-mercaptoethanol. Two stainless beads (5-mm external diameter) were added to the mixture. The tube was capped, attached to an μT-12 Beads Crusher (TAITEC, Saitama Japan), and vigorously shaken at 2,500 rpm for 3.5 min. Total RNA was isolated from 100-μl aliquots of the homogenates by using the RNeasy kit (Qiagen). cDNA was synthesized by using random hexamers and Multiscribe reverse transcriptase (Applied Biosystems, Foster City, CA, USA). Quantitative analysis of RNA transcripts was performed by real-time PCR with SYBR^®^ Green Premix Ex Taq™ (Takara, Tokyo, Japan). All oligonucleotide primers used for the real-time PCR are detailed in Table S3 in the supplemental material. Amplification of the *gapdh* gene was performed in each experiment. Each Ct value of the samples was standardized based on the *gapdh* Ct value (ΔCt), and each ΔCt value was normalized to that of the ΔCt value from PBS-treated control WT mice (ΔΔCt). Results are shown as the relative expression (1/2ΔΔCt).

### Immunization and challenge infections

#### Protective efficacy of heterologous AdHu5-prime/BDES-boost immunization against sporozoite challenge

BALB/c mice were intramuscularly immunized with a heterologous AdHu5-prime and BDES-boost regimen with a 3-week interval as described previously (14). Alternately, BALB/c mice were intramuscularly immunized with an AdHu5-prime and PBS-boost regimen with a 3-week interval. The 100-μl vaccine doses contained 5 × 10^7^ pfu of AdHu5-PfCSP or 1 × 10^8^ pfu of BDES-PfCSP. The mice were intravenously challenged with 1,000 PfCSP-Tc/Pb sporozoites at 24 h after the boost was administered, as described previously (44). The mice were checked for *P. berghei* blood-stage infection and evaluated for 1% parasitaemia as described above. Protected mice were intravenously re-challenged with 1,000 PfCSP-Tc/Pb sporozoites, and protection was defined as described above.

#### Statistical analysis

Information on the study outline, sample size, and statistical analysis is shown in the main text, figures, and figure legends. A two-tailed Fisher’s exact probability test was performed to determine the significance of differences in the protective efficacies of the vaccines, using SPSS software (version 19, Chicago, IL, USA). In all other experiments, statistical differences between the experimental groups were analysed by the methods described in the individual figure legends; *p* values of <0.05 were considered statistically significant. Statistical analyses were performed with either Prism version 7.0a (GraphPad Software Inc., La Jolla, CA, USA) or Microsoft^®^ Excel (Radmond, WA, USA).

### SUPPLEMENTAL MATERIAL

Supplemental material for this article may be found at https://XXXXXX.

## SUPPLEMENTAL FILE 1, PDF file, XXX MB

## ACKNOWLEDMENTS

TLR9^−/−^ mice were kindly provided by S. Akira (Osaka University). We would like to thank K. Takagi, K. Genshi, M. Tokutake and C. Seki, for *An. stephensi* production and animal care and T. Yoshii and T. Amano for cytokine ELISA and IVIS experiments. We are grateful to Biolegend Inc. for providing antisera. We thank Katie Oakley, PhD, from Edanz Group (www.edanzediting.com/ac) for editing the English text of a draft of this manuscript.

This work was supported, in part, by a Grant-in-Aid for Young Scientists (B) (JSPS KAKENHI grant number 26860278), the Japan Foundation for Pediatric Research (2015), and Cooperative Research Grants from NEKKEN 2014-7 (grant numbers 26-6, 27–5, 28-6 and 29-3) to M.I.; by Grants-in-Aid for Scientific Research (B) (JSPS KAKENHI grant numbers 21390126 and 25305007) and a Grant-in-Aid for Challenging Exploratory Research (JSPS KAKENHI grant number 24659460) to S.Y. T.B.E. was supported by a MEXT fellowship (153343). S.Y. and M.I. are named inventors on a patent pending related to baculovirus as a new agent against pre-erythrocytic malaria parasite (2018-57311). Neither of the products in this patent has been commercialized. All other authors declare that they have no competing interests. None of the authors have undertaken any consultancies relevant to this study.

